# NeuralRNN: a unified framework for recurrent neural network methods in cognitive neuroscience

**DOI:** 10.64898/2026.09.14.751453

**Authors:** Honghua Chen, Nai Ding

## Abstract

Recurrent neural networks (RNNs) provide a central tool in neuroscience to optimize cognitive tasks and reconstruct dynamical systems. Around these two paradigms, many RNN model variants have been tailored to capture specific computations of the brain. In practice, however, this methodological diversity remains difficult to exploit, because different paradigms and model variants are mostly implemented in idiosyncratic, study-specific code. Here we introduce NeuralRNN, an open-source Python framework that unifies these paradigms and model variants in a general-purpose pipeline. NeuralRNN brings task optimization, dynamical system reconstruction, and biological constraints into a single objective interface. It also provides an integrated and extensible model library spanning generic, structure-constrained, gain-modulated, and gated RNNs. Furthermore, data, models, and training are constructed through a shared automated pipeline and are equipped with a suite of tools for dynamical systems analysis. Together, NeuralRNN lowers the barrier to implementing, comparing, and extending RNN methods in neuroscience.

## Introduction

Cognition arises in recurrent cortical circuits, whose activity reflects both external input and the internal evolution of network state. Recurrent neural networks (RNNs) are therefore a natural model class for studying neural computation [1,2]. Since Hopfield showed that recurrent connectivity can store memories as attractors [3], RNNs have been central to theories of how circuits compute, and trained RNNs now serve as testable mechanistic hypotheses about the brain [4]. They suit this role because they balance expressivity with interpretability. Their nonlinear dynamics are rich enough to capture the state-dependent computations of cognition, yet structured enough to be dissected with dynamical systems tools [5].

Work in this tradition follows two complementary paradigms. In task optimization, an RNN is trained on a well-defined cognitive task, either by supervised learning from target outputs or by reinforcement learning from reward. The trained network is then treated as a candidate cognitive model and dissected with dynamical systems tools, which reduce its strategy to attractors, saddle points, and manifolds and reveal how the task is computed [6]. This paradigm has illuminated neural mechanisms in sensory perception, decision-making, motor control, and multi-task organization [7–9]. In the second paradigm, dynamical system reconstruction (DSR), generative RNNs are instead fitted to recorded neural or behavioral time series, so that the network’s intrinsic dynamics reproduce the observed structure and reveal how that structure arises [10–13]. Although the two paradigms ask different questions, they share the same practical needs, namely datasets, configurable models, composable training objectives, and post-training dynamical analysis.

Beyond the training paradigm, modeling brain computation requires choosing the structure of the network itself, because connectivity determines which computations a network can support and how its dynamics map onto biology [14]. Some architectures add biological structure. Excitatory-inhibitory separation enforces Dale’s law [15], multi-region architectures and spatial embedding expose cross-area routing and modularity [16,17], and short-term synaptic plasticity and gain modulation reshape dynamics beyond what connectivity alone provides [18–21]. Other architectures support analytic tractability. For example, low-rank connectivity confines dynamics to an interpretable subspace [22–24], and latent-circuit inference distills heterogeneous neural responses into a small interpretable circuit [25].

The supporting software ecosystem, however, has lagged behind this methodological diversity. Most studies are built on custom code written around one model and one task, which makes published work costly to reproduce, compare, and extend. Existing tools each cover part of the landscape, including task-optimization frameworks such as PyCog [15] and PsychRNN [26] and general-purpose brain simulators such as BrainPy [27]. What is missing is a framework in which these paradigms and architectures coexist under one interface, so that a researcher can move between them without rebuilding infrastructure.

NeuralRNN fills this gap with a unified pipeline for RNN methods in neuroscience (Figure 1). It makes three contributions. First, it provides an integrated objective system in which task optimization with supervised or reward-based signals, dynamical system reconstruction, and physiological constraints share one objective interface. Second, it establishes a model library that covers the most commonly used RNN families under a general configuration vocabulary. Third, it constructs an automated and extensible pipeline in which datasets, environments, models, and objectives are built declaratively from short configurations, while new components can be added through a public registration mechanism. Additionally, NeuralRNN also provides a suite of model-agnostic analysis tools that support fixed-point, dimensionality-reduction, and connectivity analyses of any network.

**Figure 1.**
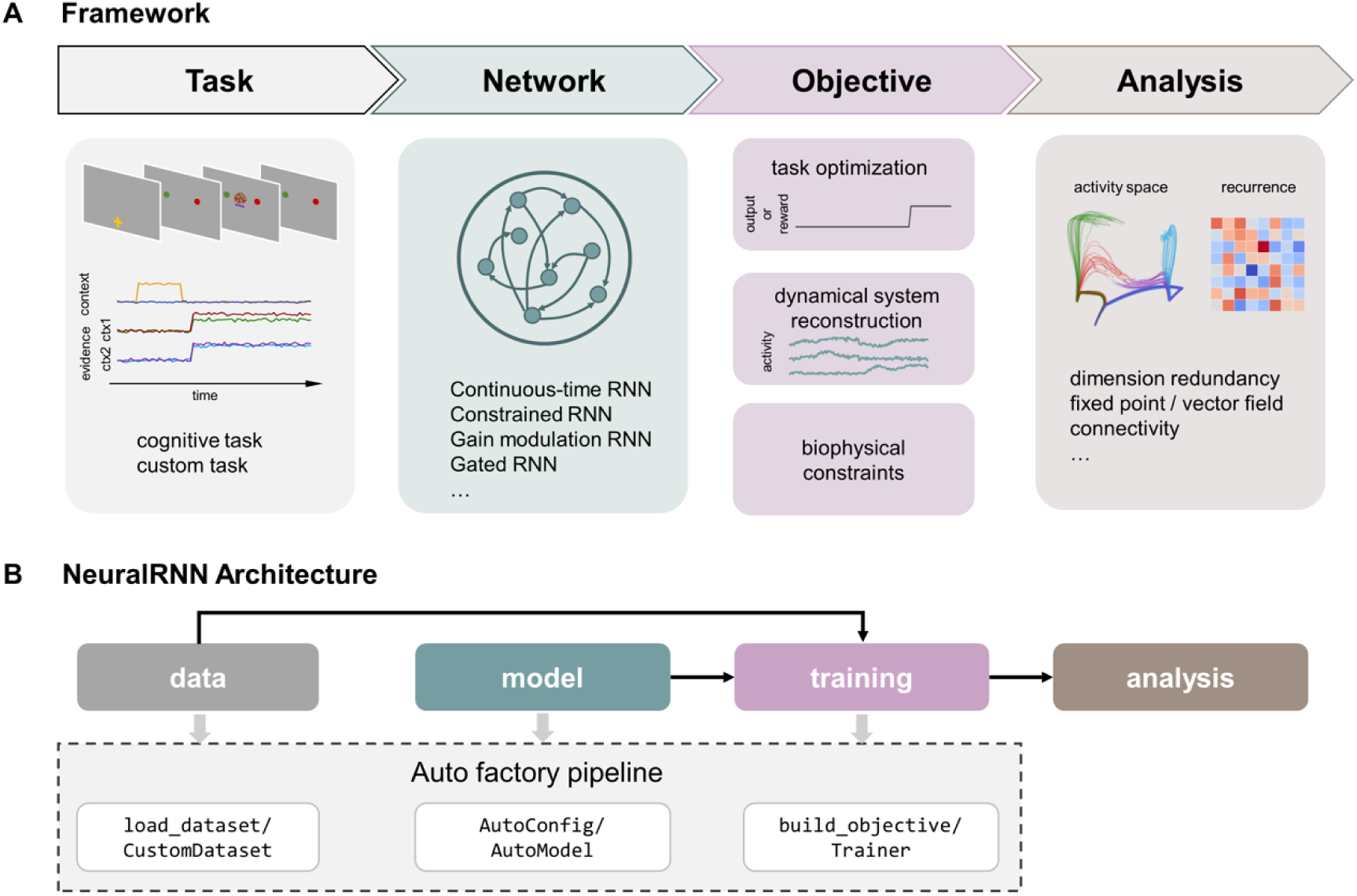
Overview of the NeuralRNN framework and software architecture. (A) Conceptual flow. Cognitive tasks provide input to RNNs, and the network layer provides configurable RNN models. The objective layer covers task optimization, dynamical system reconstruction, and biophysical constraints under one interface to train the models. The analysis layer performs model-agnostic analyses of trained networks. (B) Software architecture with four functional layers (data, model, training, analysis) served by automated factories.

## Design and implementation

### A unified framework

A researcher modeling brain computation with RNNs typically needs to do two things. One is to train a network to perform a cognitive task under supervision, or to act in an environment that rewards desired behavior. The other is to fit a network to reconstruct the recorded neural or behavioral time series, so that the model’s own dynamics explain the observations. NeuralRNN is organized around the observation that these needs are two ways of constraining the recurrent dynamics of a network. A task-optimized network is constrained through the behavior it produces, and a reconstructed network is further constrained through the activity it generates. In addition, biological knowledge also constrains the RNN model through either its activity or structure. NeuralRNN therefore implements both paradigms as different training signals applied to one model class (Figure 1A). The result is a single pipeline that runs from data to model to training to analysis, realized in four code layers (Figure 1B).

The data and training layers are where the two paradigms meet. On the data side, for task optimization, one records the inputs together with the target outputs. For reconstruction, one records the inputs together with the target system’s internal activity or external behavior. NeuralRNN gives all such datasets the same standardized batch format, so a project can freely choose which parts the network should fit, or fit several parts at once. On the training side, each choice of constraint is packaged as an objective that computes a scalar loss from a model and a batch, and a single Trainer optimizes any objective on any model. Supervised, reward-based, and reconstruction objectives, together with biological regularizers, thus become interchangeable and composable terms of one loss rather than separate training pipelines.

The model layer is designed to serve modeling questions that call for very different architectures, which are detailed in the section on flexible model construction below. These architectures differ in their equations, but they all play the same role as a recurrent dynamical system that transforms an input stream into hidden activity, together with a readout that maps activity to outputs. With input *x_t_* ∈ ℝ*^K^*, hidden state *z_t_* ∈ ℝ*^M^*, and output *y_t_* ∈ ℝ*^O^*, a model is specified as,

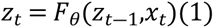

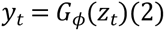

with transition parameters *θ* and readout parameters *φ*, where continuous-time models enter Eq. (1) through a common Euler discretization. This shared interface makes models interchangeable, so that any model implementing the two steps can plug unchanged into the same training and analysis suite. The same shared structure also makes the analysis layer apply to every model because analysis tools are written against the common interface of Eqs. (1)-(2) and are therefore model-agnostic. Together, the four layers form a complete pipeline from data loading to model analysis, with the details of each layer described in the following sections.

### The objective layer: task optimization, reconstruction, and biological constraints

The two paradigms differ in where their teaching signal comes from, so NeuralRNN packages every training signal as an objective, a configurable module that computes a scalar loss from a model and a batch of data. The `Trainer` therefore calls only `compute_loss(model, batch)` and remains agnostic to where the loss comes from. Figure 2 shows where these signals act on a network. Three families of built-in objectives cover the paradigms and their combination.

**Figure 2.**
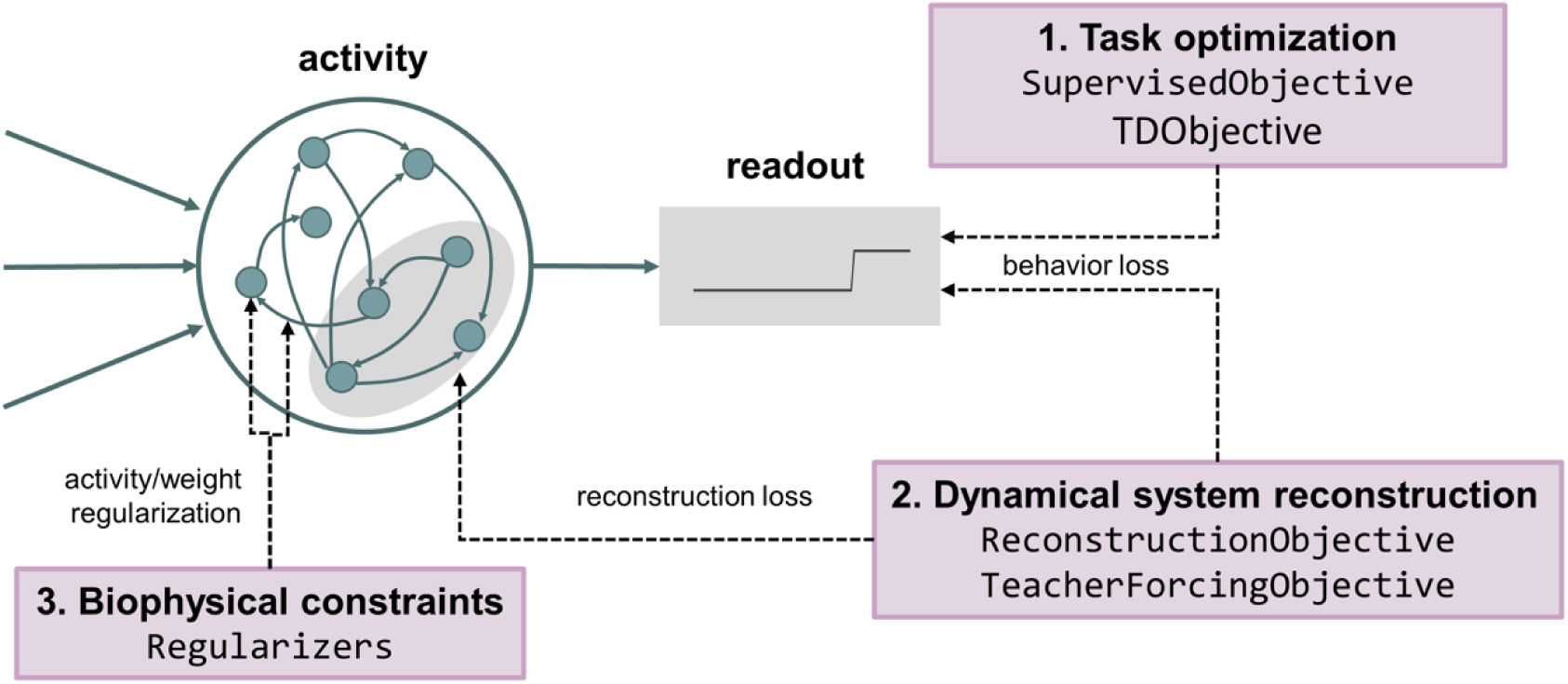
Unified objective abstraction. Dashed arrows mark where the composable training signals act. These are (1) task optimization, with losses acting on the model readout, (2) dynamical system reconstruction, which constrains internal activity to reproduce recorded dynamics and optional model readout, and (3) biophysical constraints, which regularize network activity and weights.

#### Task optimization

Supervised training applies when the desired behavior can be written down as target outputs. `SupervisedObjective` computes a behavioral loss on the readout output (cross-entropy for classification, masked mean-squared error for regression). Many behaviors of interest, however, have no target outputs. In navigation, foraging, or value-based decision-making, the correct action sequence is unknown and only reward feedback is available, so the network must discover its strategy through interaction. For these problems, policy-gradient losses (advantage actor-critic [28] and proximal policy optimization (PPO) [29]) train agents through live environment interaction, and `TDObjective` trains a critic by semi-gradient temporal-difference learning on recorded episodes [30].

#### Reconstruction

`ReconstructionObjective` fits teacher data as a weighted sum of a behavioral term and an activity term. Three switches adapt it to different reconstruction settings. For example, `state_map` inserts a learnable embedding when the model and the recorded activity live in spaces of different dimensionality, `recorded_dims` restricts the fit to a subset of recorded units, and `activity_fn` selects the nonlinearity applied to the model’s activity before comparison. Teacher-student network reconstruction [23] and low-dimensional circuit embedding [25] are thereby covered by a single class. `TeacherForcingObjective` implements generalized teacher forcing (GTF) [31], which stabilizes long-sequence training on chaotic systems by sparsely forcing *z* = *αz*_obs_ +(1 ― *α*) *z*_pred_, with the forcing strength *α* annealed by the Trainer.

#### Biological constraints

Beyond the paradigm objectives, NeuralRNN provides a library of regularizers, including activity-L2, weight-L2, and input-output orthogonality penalties, which are exposed as reusable components that can be attached to any objective. For common use cases, NeuralRNN has ready-made objectives to bundle them. For example, `RegularizedSupervisedObjective` adds activity and weight penalties to the task loss, and `ConstrainedSupervisedObjective` adds structural penalties reported by the model itself, such as the wiring-cost term of the spatially embedded model [17].

Together, all objectives share a single composition rule,

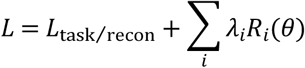

With *L*_task/recon_ for the task or reconstruction loss, *R_i_* for regularization or structural terms, and *λ_i_* for their coefficients. Each term can be enabled and weighted independently, so behavioral, dynamical, and structural constraints can be combined within one training run. This combination is what joint modeling of task behavior and neural activity requires in practice.

### Flexible model construction

Different modeling questions demand different network structures, and the model layer reflects this. It registers 17 model types, organized into families by the modeling problem they address (see S1 Appendix for more detailed model descriptions):

1. *Continuous-time rate models*. The default hypothesis in cognitive modeling is a circuit of units evolving in continuous time, whose trained dynamics can be read as a candidate mechanism for the circuit under study. The `ctrnn` model class provides this model and serves as the base architecture from which the families below can inherit (Fig. 3A).
2. *Structure-constrained models*. Cortical connectivity in the real brain is not generic. Neurons obey Dale’s law, wiring is sparse and modular, connections cluster within and between areas, and much of the variance of population activity is low-dimensional. Building such structure into a model lets the researcher ask which constraints a computation actually requires (Fig. 3B-E). The `constrained_rnn` model provides hard masks on the recurrent, input, and output pathways as the general mechanism, and the more specific architectures inherit from it. Based on the `sparse_rnn` and `modular_rnn` that hard-wire sparse and modular topologies, the `multiarea_rnn` implements multi-area networks as the special case of block-structured connectivity masks (Fig. 3B) [16]. The `se_rnn` ports the spatially embedded RNN, in which neurons occupy positions in space and wiring cost enters the loss (Fig. 3E) [17]. Two further models constrain the network in other forms. `ei_rnn` enforces Dale’s law by splitting neurons into excitatory and inhibitory populations with sign-constrained outgoing weights (Fig. 3C) [15], and `lowrank_rnn` parameterizes *W*_rec_ = *mn*^T^/*N* (*m*,*n* ∈ ℝ*^N^*^×*R*^, rank *R*), confining the dynamics to an interpretable low-dimensional subspace (Fig. 3D) [22–24].
3. *Gain-modulated and plasticity models*. Neural computation is shaped not only by connectivity but also by neuromodulation and synaptic dynamics, which act on the activation function rather than the weight matrix. `gain_rnn` adds per-neuron gain and bias to the firing-rate map, with `gain_position` selecting output gain (amplitude) or input gain (slope, corresponding to targeted gain modulation [19]) (Fig. 3F). `stp_rnn` treats Tsodyks-Markram short-term plasticity as a dynamic gain with effective efficacy syn_*x* ⋅ syn_*u* (Fig. 3G), ported from synaptic theories of working memory [18,20].
4. *Gated architectures*. Some modeling goals call for the architectures standard in machine learning, such as tasks with long temporal dependencies, behavioral-strategy discovery from animal choice data [13], or direct comparison with the machine-learning literature. `gated_rnn` provides GRU [32] and LSTM [33], with the gate equations implemented as an explicit per-step recurrence (Fig. 3H) so that the fixed-point and linearization tools of the analysis layer apply unchanged.
5. *Models for reconstruction*. Reconstructing dynamics from data places its own demands on the model. The reconstructed system should admit analytic inspection of its fixed points and Jacobians, and heterogeneous high-dimensional recordings should be distillable into an interpretable circuit. The piecewise-linear RNN family for DSR (shallow PLRNN [12], dendritic dendPLRNN [34], and almost-linear RNN [35]) replaces the smooth nonlinearity with piecewise-linear maps. This exposes analytic Jacobians and fixed points and drives the analytic backend of the analysis layer. The latent-circuit model [25] embeds a low-dimensional recurrence into neural space through an embedding matrix (Fig. 3I), so that a high-dimensional trained or biological network can be distilled into a small circuit that recovers its essential dynamics.
6. *Actor-critic agents*. Sequential decision problems such as navigation, foraging, and planning cannot be trained by supervision, because no target action sequence exists. The agent must act, observe reward, and improve its policy. The actor-critic model turns any registered RNN core into an RL agent by attaching policy and value readout heads (Fig. 3J). The policy head produces a categorical distribution over discrete actions or a diagonal Gaussian over continuous actions, and it can be omitted entirely for critic-only value agents [30]. An optional auxiliary head predicts environment variables from the hidden state, serving as a learned world model for planning agents [36]. Because the agent is itself a NeuralDynamicsModel, RL-trained networks accept the same analysis tools as any other model.

**Figure 3.**
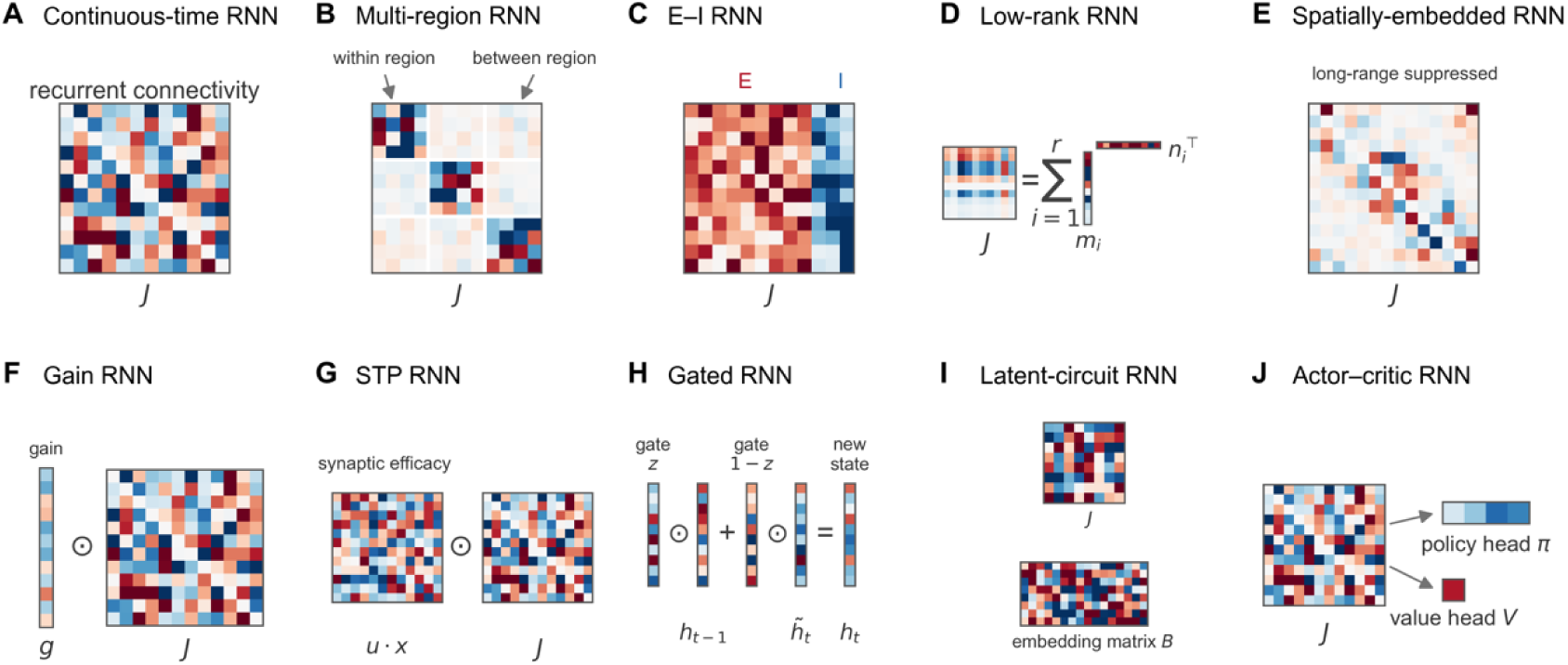
Built-in models in NeuralRNN. (A) Continuous-time RNN, the default model with an unstructured recurrent connectivity *J*. (B) Multi-region RNN, whose block-structured connectivity separates strong within-region from sparser between-region connections. (C) E-I RNN, which obeys Dale’s law: each column of *J* is constrained to be either excitatory or inhibitory. (D) Low-rank RNN, in which *J* is parameterized as a sum of *r* rank-one terms m_i_ n ^T^, confining the network dynamics to a low-dimensional subspace. (E) Spatially-embedded RNN, in which long-range connections are suppressed. (F) Gain RNN, which modulates the recurrent computation through a per-neuron gain vector *g* applied element-wise to the connectivity. (G) STP RNN, in which short-term plasticity gives each synapse a dynamic efficacy *u·x* that rescales the effective connectivity from moment to moment. (H) Gated RNN, such as GRU or LSTM, in which the state update mixes the previous state *h_t-1_* with a candidate state *h̃_t_* through learned gates *z* and *1−z*. (I) Latent-circuit RNN, a low-dimensional recurrence whose activity is mapped into neural space through an embedding matrix *B*, so that it can be fitted directly to recorded neural activity. (J) Actor-critic RNN, in which any recurrent core is equipped with a policy head *π* producing action distributions and a value head *V* estimating expected returns for reinforcement learning. Red and blue denote positive and negative weights, respectively. The full list of 17 model types, with equations and configuration options, is given in S1 Appendix.

Across all families, there is a set of shared configuration options. All models share the Euler update *z*_′_ = (1 ― *α*)*z* + *αf*(⋅), with the update fraction *α*. The option `nonlinearity_mode` specifies where the nonlinearity sits within the Euler step. Published studies differ in this detail, which is easy to overlook but changes the effective dynamics. Four `freeze_*` flags provide a uniform vocabulary for freezing input weights, recurrent weights, the initial state, and output weights, making reservoir-style training and continual learning straightforward. In addition, the shape of the nonlinearity (e.g., ReLU, Sigmoid, and tanh) and the dimensions of the input, recurrent, and output pathways are likewise configurable in every family.

### The automated pipeline

Assembling a project from these diverse components requires configuration, and NeuralRNN therefore adapts the configuration-registry-automation pattern of Transformers [37], giving each layer an automated factory (Figure 1B).

#### Datasets

Modeling starts from one of three kinds of data sources, namely a task specification for supervised optimization, a set of recorded arrays for reconstruction, or an interactive environment for reward-based optimization. The data layer is based on a single contract, `BaseDataset`, whose `sample_batch()` method returns a standardized batch dictionary (inputs, targets, mask, activity, and so on) for training. Task specifications are served by built-in cognitive-task classes covering decision-making, perception, and working memory, together with the multi-task suites of Yang et al. [9] and Driscoll et al. [38]. An optional `NeurogymDataset` adapter gives access to the NeuroGym task library [39], and new tasks can be specified through `task_base` by implementing a `generate_trials()` interface that defines inputs and outputs. For reconstruction, recorded data enter through `ReconstructionDataset`, which imports time series from arrays or files. For reinforcement learning, the data source is not a fixed batch but an interactive environment. Environments follow a Gymnasium-style contract and are vectorized through `SyncVectorEnv`. Under this contract, `reset_env` returns an initial observation and `step_env` returns the next observation, reward, termination flags, and an info dictionary that carries task variables for later neural analysis. For offline value learning, recorded episodes can instead be loaded as ordinary datasets, without an environment loop. Lastly, DATASET_REGISTRY and `load_dataset()` give name-based access to all registered datasets, including user-registered ones.

#### Models

Configurations derive from `NeuralRNNConfig`, each declaring a model_type key and serializable to JSON. Models derive from `NeuralDynamicsModel` and implement a `save_pretrained()`/`from_pretrained()` contract for checkpoints. The @register_model and @register_config decorators add user components to the registries, after which `AutoConfig.for_model(name, overrides)` and `AutoModel.from_config(config)` construct them by name.

#### Objectives and training

Objectives have their own registry and factory. `build_objective(name, kwargs)` constructs any objective from the previous section, and @register_objective supports user extensions. The Trainer is a single paradigm-agnostic loop in which each step draws a batch from `sample_batch()` and calls `compute_loss(model, batch)` to obtain and backpropagate the scalar loss.

`TrainingArguments` configures gradient clipping, optimization, logging, evaluation, checkpointing, hidden-state dropout, and early stopping. For reinforcement learning, `RLTrainer` mirrors the same design for on-policy learning. Each update collects a fixed number of steps from the environments under the current policy and estimates returns and advantages with n-step or generalized advantage estimation [40]. The recorded sequences are then replayed through the recurrent network for backpropagation through time, with hidden states reset at episode boundaries so that temporal structure is preserved. Periodic evaluation, early stopping, and checkpointing follow the same conventions as the Trainer.

### Analysis tools

Trained networks are analyzed with a model-agnostic tool suite. Every tool depends only on the public interface of `NeuralDynamicsModel` (recurrence, readout, jacobian, generate) and never imports a concrete model class, so any conforming model, including user-registered ones and RL agents, automatically gains the full suite. At the activity level, the tools characterize the dynamical system directly through fixed-point analysis with linearization, stability analysis, and vector-field sampling, and through dimensionality reduction including PCA, demixed PCA [41], and CCA-based alignment [42]. At the behavioral level, psychometric-curve tools quantify task performance. A separate visualization layer provides static and animated displays, including trajectory plots, fixed points and vector fields, and psychometric curves.

## Results

We first illustrate the breadth of RNN modeling the framework covers and then show usage examples that demonstrate its usability and extensibility.

### A framework with broad coverage

Table 1 compares NeuralRNN with five existing tools for RNN modeling in neuroscience along the paradigms and the RNN architectures. NeuralRNN is, to our knowledge, the only framework that combines task optimization and dynamical system reconstruction, and provides the architectures listed above.

**Table 1:** Feature coverage of NeuralRNN and existing tools.

| Feature | NeuralRNN | PyCog<br>[15] | PsychRNN<br>[26] | PyRates<br>[43] | nn4n<br>[44] | BrainPy<br>[27] |
| --- | --- | --- | --- | --- | --- | --- |
| **Modeling paradigms** |  |  |  |  |  |  |
| Task optimization (supervised) | Yes | Yes | Yes | Yes | Yes | Yes |
| Task optimization (reinforcement learning) | Yes | No | No | No | No | No |
| Dynamical system reconstruction | Yes | No | No | No | No | No |
| **RNN architectures** |  |  |  |  |  |  |
| Continuous-time RNN | Yes | Yes | Yes | Yes | Yes | Yes |
| Constrained connectivity | Yes | No | Partial | No | Partial | No |
| Dale excitatory-inhibitory constraints | Yes | Yes | Yes | No | Yes | Yes |
| Low-rank connectivity | Yes | No | No | No | No | No |
| Gain modulation | Yes | No | No | No | No | Yes |
| Gated architectures (GRU and LSTM) | Yes | No | Partial | No | No | Yes |
| Actor-critic | Yes | No | No | No | No | No |

To evaluate the framework, we used NeuralRNN to reproduce the modeling and analysis workflows of a broad set of published studies (see S1 Appendix), covering supervised [6,24,38,45–48] and reward-based [30,36,49–51] task optimization as well as dynamical system reconstruction [13,23,25,52]. These studies span domains including sensory perception, decision-making, working memory, motor control, navigation, planning, learning, and multi-task organization, indicating the broad usage of NeuralRNN for general-purpose modeling of the brain. Complete reproduction details are provided in the online documentation at https://neuralRNN.readthedocs.io/.

Notably, the unified framework in NeuralRNN is more than a programming convenience. The two paradigms constrain the same brain from different sides, providing different perspectives for modeling the brain. A model of the brain should both do what the brain does and vary the way neural activity varies, under the structural constraints that biology imposes. Writing behavioral, dynamical, and structural constraints in terms of one objective makes their combination straightforward, and several recent modeling advances are precisely built on this fusion. For example, latent-circuit inference jointly optimizes over behavioral and neural activity data to infer a latent circuit, providing a robust explanation of the underlying network dynamics [25]. Similarly, connectome-constrained networks are trained on a task with partially-fixed connectivity to investigate how far the network remains identifiable under such constraints [52].

NeuralRNN also facilitates direct comparison across models and tasks. Networks trained on the same task with different architectures can reach comparable performance while relying on markedly different internal dynamics [53], and the shared pipeline of NeuralRNN makes this sensitivity measurable rather than anecdotal.

### Usage of NeuralRNN: from a cognitive task to a reconstructed circuit

We show a complete project that uses both paradigms on a context-dependent decision-making task (Figure 4 and Listing 1) [6]. In the first paradigm, task optimization, a continuous-time RNN is trained on this task, in which a context cue instructs the network to report either the motion or the color evidence of a random-dot stimulus (Figure 4A). The dataset is generated from the built-in task registry, the model is built from a short configuration, and a regularized supervised objective trains it with the generic Trainer. The trained network performs the task with context-appropriate psychometric curves, and its low-dimensional trajectories separate first by context and then by accumulated evidence, reproducing the characteristic geometry reported for this task.

**Figure 4.**
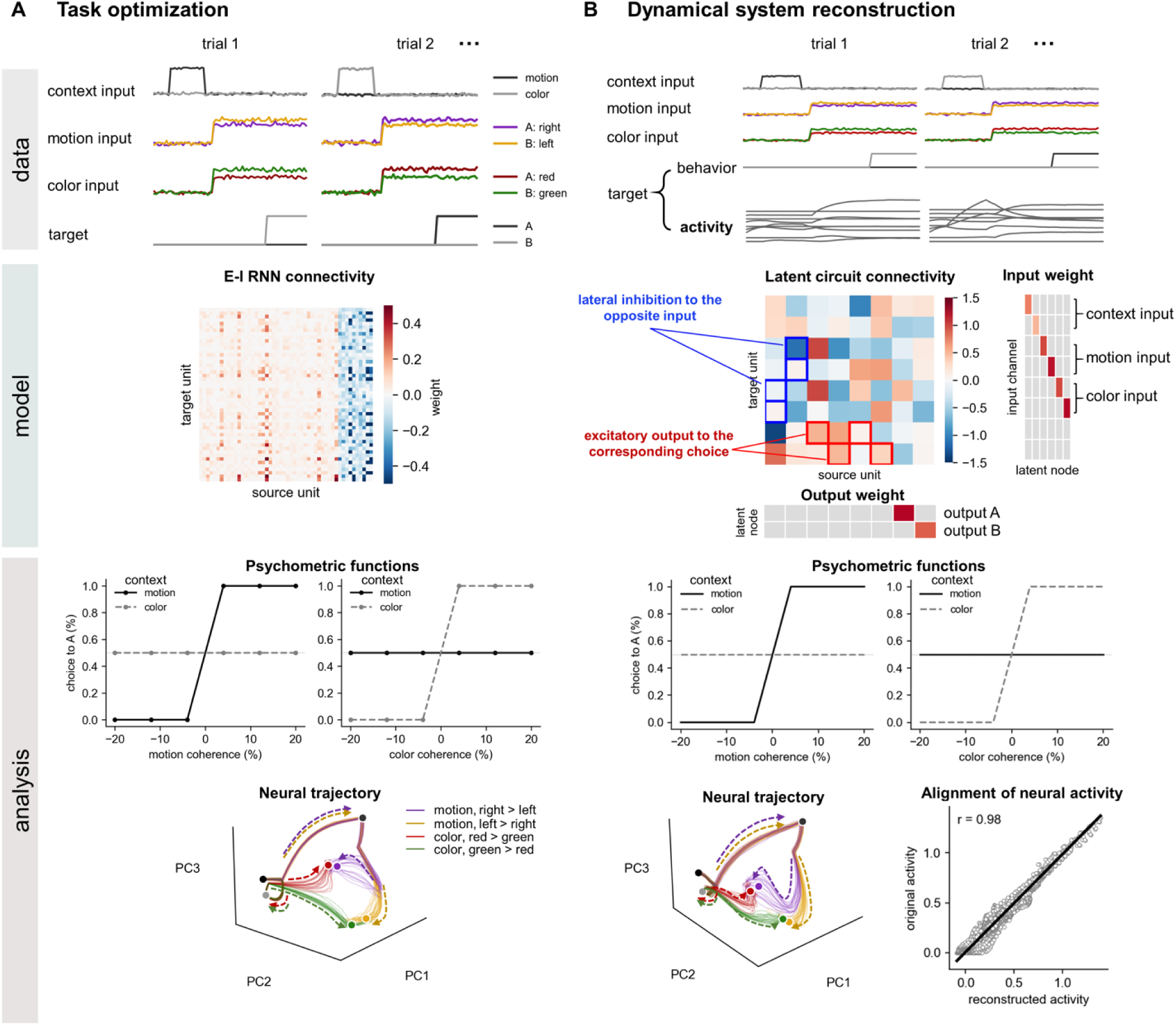
Usage of two modeling paradigms. (A) Task optimization. The data section shows example trials of the context-dependent decision-making task, in which a context cue instructs the network to report either the motion or the color evidence, and the target is the correct behavioral response. The model section shows the recurrent connectivity of the trained E-I RNN. The analysis section shows context-appropriate psychometric functions, with choice depending only on the cued evidence stream, and low-dimensional neural trajectories that separate first by context and then by accumulated evidence. (B) Dynamical system reconstruction. The data section shows the same task, but the training target now comprises recorded behavior and neural activity rather than a supervised output. The model section shows the inferred latent circuit with the input and output weights. The connectivity reveals an interpretable mechanism in which units driven by contexts inhibit the units driven by opposite evidence laterally, while units driven by evidence send excitatory output to the corresponding choice. The analysis section shows that the reconstructed circuit reproduces the psychometric functions and the geometry of the neural trajectories of the source network, and that its reconstructed activity quantitatively matches the original activity (r = 0.98).

In the second paradigm, the trained network becomes the data source (Figure 4B). `ReconstructionDataset` records its activity with noise on the same task, and a latent-circuit model [25] with eight latent units is trained to reconstruct that activity through the reconstruction objective, with an orthogonality constraint maintained after each gradient step. The inferred circuit reproduces the task behavior, and its embedded states quantitatively match the original network’s trajectories. Importantly, the connectivity reveals an interpretable mechanism in which units driven by contexts use inhibitory connections to suppress the units driven by evidence in the opposite dimension, while units driven by evidence send excitatory output to the corresponding choice. Here, a trained network serves as the reconstruction target for demonstration, but the same workflow applies directly to recorded neural data. Reconstructing a biological circuit therefore requires no change beyond the data source. Because the two paradigms share the dataset contract, the objective interface, and the Trainer, moving from training a network to reverse-engineering it changes only the dataset, the model type, and the objective, each still a one-line factory call.

**Listing 1.**
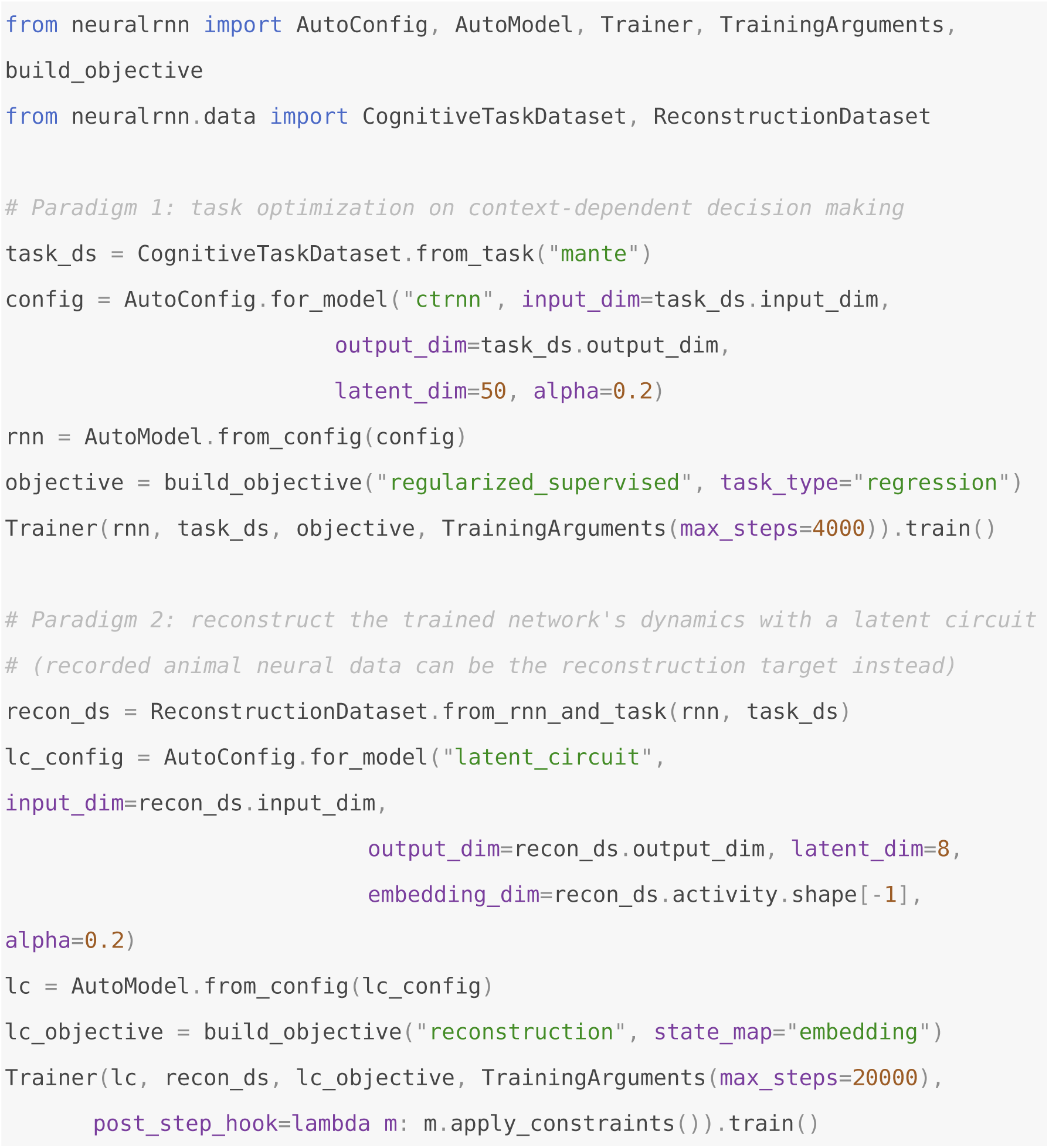
A complete two-paradigm project.

### Building new components

The framework is designed to be extended with new components, which follow the same contract as the built-ins. A new model subclasses `NeuralDynamicsModel`, implements `recurrence()` and `readout()`, and registers a name. The base class then supplies state initialization, rollout, free-running generation, parameter freezing, checkpointing, and access to every analysis tool. A new objective subclasses Objective, implements `compute_loss(),` and registers a name, after which it can compose with each model class with the Trainer. Listing 2 shows both additions in miniature. The same pattern extends to reinforcement learning, where new environments and new policy-gradient losses register through the equivalent decorators. Lastly, the framework’s built-in model types also serve as worked examples of the extension process, ranging from single-file models to multi-file families.

**Listing 2.**
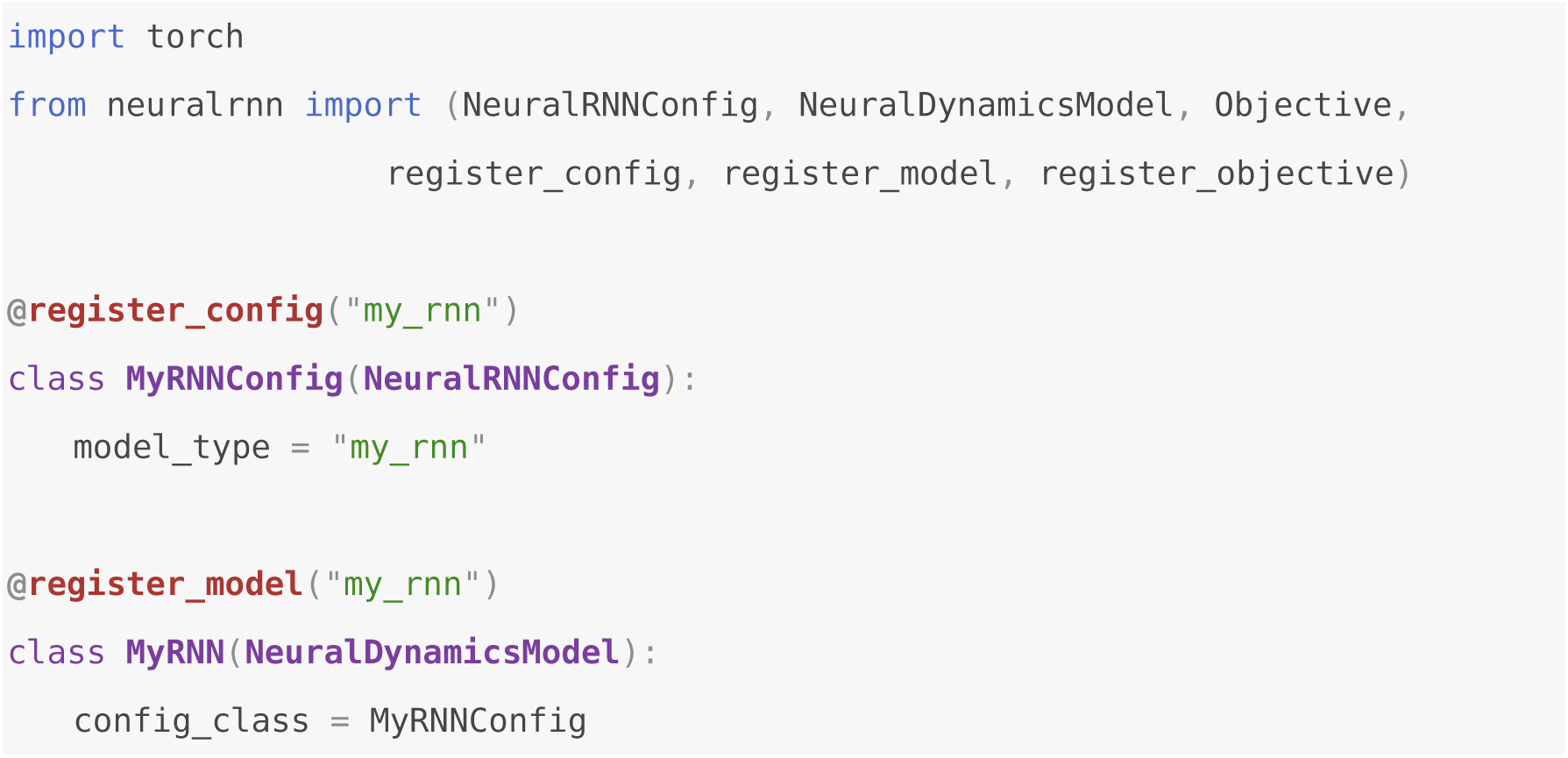
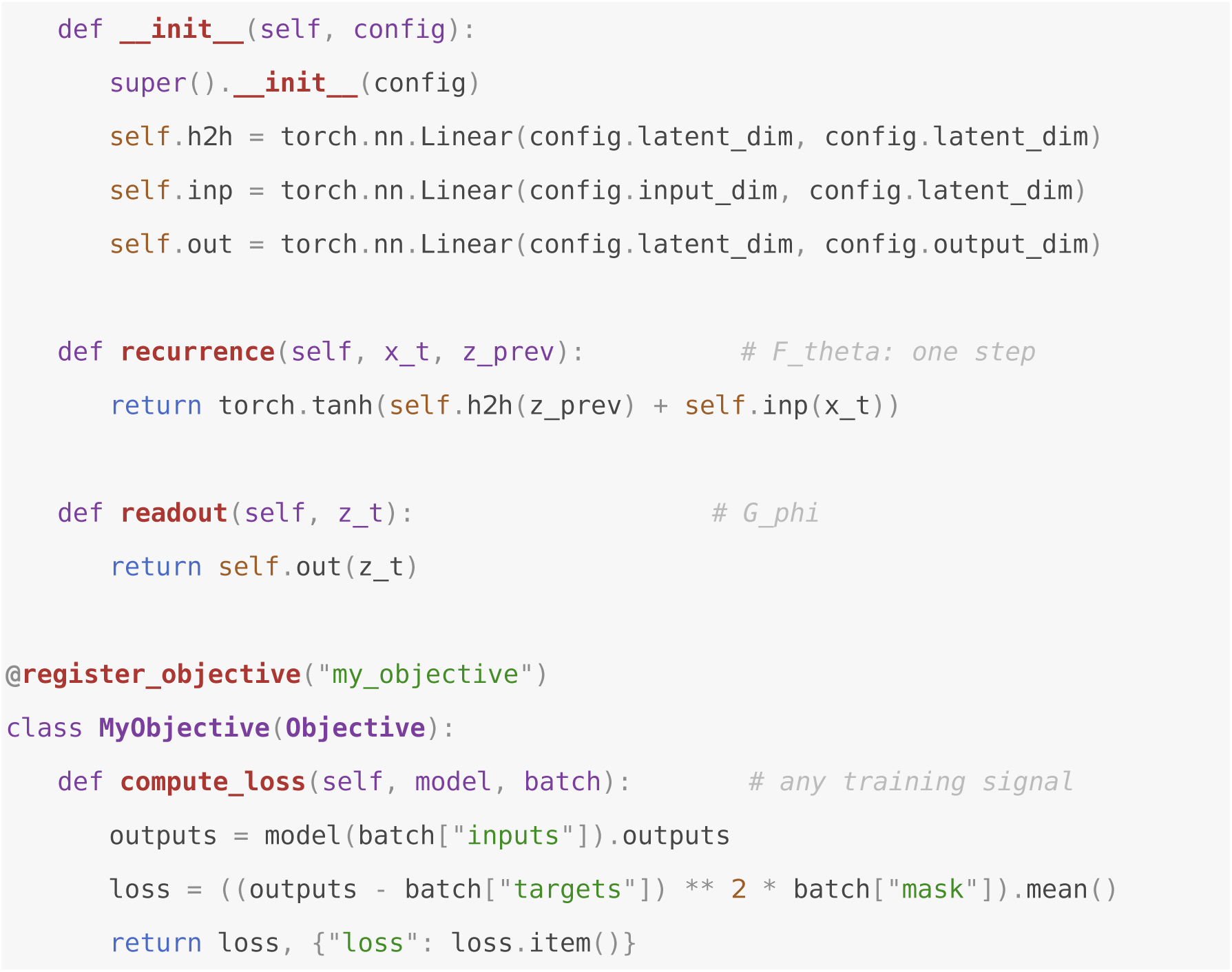
Extending the framework with a new model and a new objective.

## Availability and future directions

### Availability

NeuralRNN is released under the MIT license at https://github.com/ChenHH730/NeuralRNN, with online documentation at https://neuralRNN.readthedocs.io/. The framework is written in Python and supports Python 3.9 and above. Its only hard dependencies are PyTorch (version 2.0 or above), NumPy, SciPy, safetensors, Matplotlib, and tqdm. The NeuroGym task adapter and the reinforcement-learning environments additionally use NeuroGym and Gymnasium, which are loaded on demand. All code is enforced by an automated test suite that is shipped with the package and runs with pytest.

### Limitations

The framework covers rate-based and gated recurrent networks in discrete time. Because the unified abstraction is built on a discrete-time single-step transition map, it does not address spiking networks, multi-compartment biophysical models, or stochastic and delay differential equations, although any model that can be written in the form of Eqs. (1)-(2) can be added through the same interface. On the reinforcement-learning side, the framework currently implements policy-gradient algorithms (advantage actor-critic and PPO) and temporal-difference value learning.

### Future directions

Two directions follow naturally from the current design. First, the training layer currently assumes gradient-based optimization through time. Adding dedicated initialization schemes and FORCE-type learning rules [54]. Second, the registry mechanism leaves room for community-contributed tasks, environments, models, and analyses.

